# Mutations in the *Drosophila* splicing regulator *Prp31* as a model for Retinitis pigmentosa 11

**DOI:** 10.1101/147918

**Authors:** Malte Lehmann, Sarita Hebbar, Holger Brandl, Weihua Leng, Naharajan Lakshmanaperumal, Sylke Winkler, Elisabeth Knust

## Abstract

Retinitis pigmentosa is a clinically heterogeneous disease affecting 1.6 million people worldwide. A growing number of identified disease-causing genes are associated with the spliceosome, but the molecular consequences that link defects in splicing factor genes to the aetiology of the disease remain to be elucidated. In this paper, we present a *Drosophila* model for Retinitis pigmentosa 11, a human disease caused by mutations in the splicing factor PRPF31. Here, we induced mutations in the *Drosophila* orthologue *Prp31*. Mutant flies are viable and show a normal eye phenotype when kept under regular light conditions. However, when exposed to constant light, photoreceptors of mutant flies degenerate, thus resembling the human disease phenotype. Degeneration could be shown to be associated with increased oxidative stress. This increase was in agreement with severe dysregulation of genes involved in oxidation/reduction processes, as revealed by high throughput transcriptome sequencing. In fact, light induced photoreceptor cell degeneration could be attenuated by experimentally reducing oxidative stress. A comparable decrease in retinal degeneration was achieved by raising mutant larvae on a vitamin A-depleted medium, thereby reducing rhodopsin levels. Finally, transcriptome data further uncovered an overall retention of introns in mRNAs. Among those, mRNAs of genes involved in synapse assembly, growth and stability were most prominent. These results point to a multifactorial genesis of light induced degeneration in retinae of *Prp31* mutant flies, including transcriptional and splicing dysregulation, oxidative stress and defects in vitamin A metabolism.

## Introduction

Retinitis pigmentosa (RP; OMIM 268000) is a clinically heterogeneous set of retinal dystrophies, which affects about 1.6 million people worldwide. It often starts with night blindness in early childhood due to the degeneration of rod photoreceptor cells (PRCs), continues with the loss of the peripheral visual field caused by degeneration of cones (tunnel vision), and progresses to complete blindness in later life. RP is a genetically heterogeneous disease, and can be inherited as autosomal dominant (adRP), autosomal recessive (arRP) or X-linked (xlRP) disease. So far >50 genes have been identified that are causally related to non-syndromic RP (Daiger et al., 2014) (see RetNet: http://www.sph.uth.tmc.edu/RetNet/disease.htm). Affected genes are functionally diverse. Some of them are expressed specifically in PRCs and encode, among others, transcription factors (e. g. *CRX*, an *otx*-like photoreceptor homeobox gene), components of the light-induced signalling cascade, including the visual pigment rhodopsin (Rho/*RHO* in *Drosophila*/human), or genes controlling vitamin A metabolism (e.g. *RLBP-1*, encoding Retinaldehyde-binding protein). Other genes are associated with the control of cellular homeostasis, for example *CRB1*, a gene required for the maintenance of polarity. Interestingly, the second-largest group of genes causing adRP, comprising 7 of 23 genes known, encodes regulators of the splicing machinery. So far, mutations in five PRPF (**pr**emRNA **p**rocessing **f**actor) genes, *PRPF3*, *PRPF4*, *PRPF6, PRPF8* and *PRPF31*, have been linked to adRP, namely RP18, RP70, RP60, RP13 and RP11, respectively. *PAP1* (Pim1-associated protein) and *SNRNP200* (small nuclear ribonuclearprotein-200), two other genes involved in splicing, have been suggested to be associated with RP9 and RP33, respectively (Maita et al., 2004; Zhao et al., 2009) [reviewed in (Liu and Zack, 2013; Mordes et al., 2006; Poulos et al., 2011; Ruzickova and Stanek, 2016)]. The five *PRPF* genes encode components regulating the assembly of the U4/U6.U5 tri-snRNP, a major module of the pre-mRNA spliceosome machinery (Will and Luhrmann, 2011). Several hypotheses have been put forward to explain why mutations in ubiquitously expressed components of the general splicing machinery show a dominant phenotype only in PRCs. One hypothesis suggests that PRCs with half the copy number of genes encoding general splicing components cannot cope with the elevated demand of RNA-/protein synthesis required to maintain the exceptionally high metabolic rate of PRCs in comparison to other tissues. Hence, halving their gene dose eventually results in apoptosis. Although this model is currently favoured, other mechanisms, such as impaired splicing of PRC-specific mRNAs or toxic effects caused by accumulation of mutant proteins have been discussed and cannot be excluded to contribute to the disease phenotype [discussed in (Mordes et al., 2006; Scotti and Swanson, 2016; Tanackovic et al., 2011)].

The observation that all adRP-associated genes involved in splicing are highly conserved from yeast to human allows to use model organisms to unravel the genetic and cell biological functions of these genes, which ultimately will provide a mechanistic characterization of the origin of the diseases. In the case of RP11, the disease caused by mutations in *PRPF31*, three mouse models have been generated by knock-in and knock-out approaches. Unexpectedly, mutant mice did not show any sign of retinal degeneration (Bujakowska et al., 2009). Further analyses revealed that the retinal pigment epithelium, rather than the PRCs, is the primary tissue affected in *Prpf31* heterozygous mice (Farkas et al., 2014; Graziotto et al., 2011). Morpholino-induced knock-down of zebrafish *Prpf31* results in strong defects in PRC morphogenesis and survival (Linder et al., 2011). Defects obtained by retina-specific expression of zebrafish *Prpf31* constructs that encode proteins with the same mutations as those mapped in RP11 patients (called AD5 and SP117, respectively) were explained to occur by either haplo-insufficiency or by a dominant-negative effect of the mutant protein (Yin et al., 2011). In *Drosophila*, no mutations in the orthologue *Prp31* have been identified so far, but RNAi-mediated knock-down of *Prpf31* in the developing *Drosophila* eye induced, besides strong developmental defects of the eye, signs of PRC degeneration (Ray et al., 2010)

In order to establish a meaningful *Drosophila* model for RP11-associated retinal degeneration, we isolated two mutant alleles of *Prp31*, *Prp31^P17^* and *Prp31^P18^*, which carry missense mutations that result in exchanges of conserved amino acids. Flies heterozygous for either of these mutations are viable and develop normally. Strikingly, when exposed to constant light, mutant flies undergo retinal degeneration. Degeneration of mutant PRCs was associated with increased oxidative stress. Consistent with this, transcriptome analyses from heads of *Prp31^P18^* homozygous flies showed transcriptional dysregulation of genes involved in oxidation/reduction processes. In addition, an overall retention of introns in mRNAs was observed. Retinal degeneration could be ameliorated by supplementing the food with NSC23766, a known inhibitor of NADPH (nicotinamide adenine dinucleotide phosphate)-oxidase activity, or by raising larvae on a vitamin A-depleted medium. From these results, we conclude a multifactorial genesis of *Prp31*-linked retinal degeneration in flies.

## Results

### Flies heterozygous for mutations in *Prp31* undergo light-dependent retinal degeneration

It was recently shown that RNAi-mediated knockdown of *Drosophila Prp31* in the eye using *eyeless* (*ey*)-Gal4 or GMR-Gal4 results in smaller eyes or no eyes at all and degeneration of photoreceptor cells (PRCs) and pigment cells (Ray et al., 2010). Since *eyeless* is expressed already in the early eye imaginal disc prior to PRC differentiation, some of the defects observed could be secondary, for example as a consequence of defective cell fate specification.

To establish a more meaningful *Drosophila* model for RP11-associated retinal degeneration, which would allow a deeper insight into the role of this splicing factor in the origin and progression of the disease, we set out to isolate specific mutations in *Drosophila Prp31* by TILLING (**T**argeting **I**nduced **L**ocal **L**esions **IN G**enomes), following a protocol described recently (Spannl et al., 2017). In total, 2.400 genomes of EMS (ethyl methanesulfonate)- mutagenized flies were screened for sequence variants in two different amplicons of *Prp31*. Four sequence variants were identified, which were predicted to result in potentially deleterious missense mutations. Two of the four lines were recovered from the living fly library and crossed for three generations to wild-type (*w^1118^*) flies to reduce the number of accompanying sequence variants. In two of the lines, named *Prp31^P17^* and *Prp31^P18^*, mutations in *Prp31* could be verified. *Prp31^P18^* was viable as homozygotes or in trans over *Df(3L)Exel6262*, which removes, among others, the *Prp31* locus. In contrast, no homozygous *Prp31^P17^* flies were obtained. However, *Prp31^P17^* was viable in trans over *Prp31^P18^* and over *Df(3L)Exel6262*. This suggests that the lethality was due to a second site mutation, which was not removed despite extensive out-crossing. The molecular lesions in the two alleles were mapped in the protein coding region of *Prp31*. *Drosophila* PRP31 is a protein of 501 amino acids, which contains a NOSIC domain (named after the central domain of Nop56/SIK1-like protein), a Nop (**N**ucle**o**lar **p**rotein) domain required for RNA binding, a PRP31 _C-specific domain and a nuclear localization signal, NLS. *Prp31^P17^* contained a point mutation that resulted in a non-conservative glutamine to arginine exchange (G90R) N-terminal to the NOSIC domain. *Prp31^P18^* contained a non-conservative exchange of a proline to a leucine residue in the Nop domain (P277L) (Fig. 1A and Supplementary Fig. S1).

**Figure 1:**
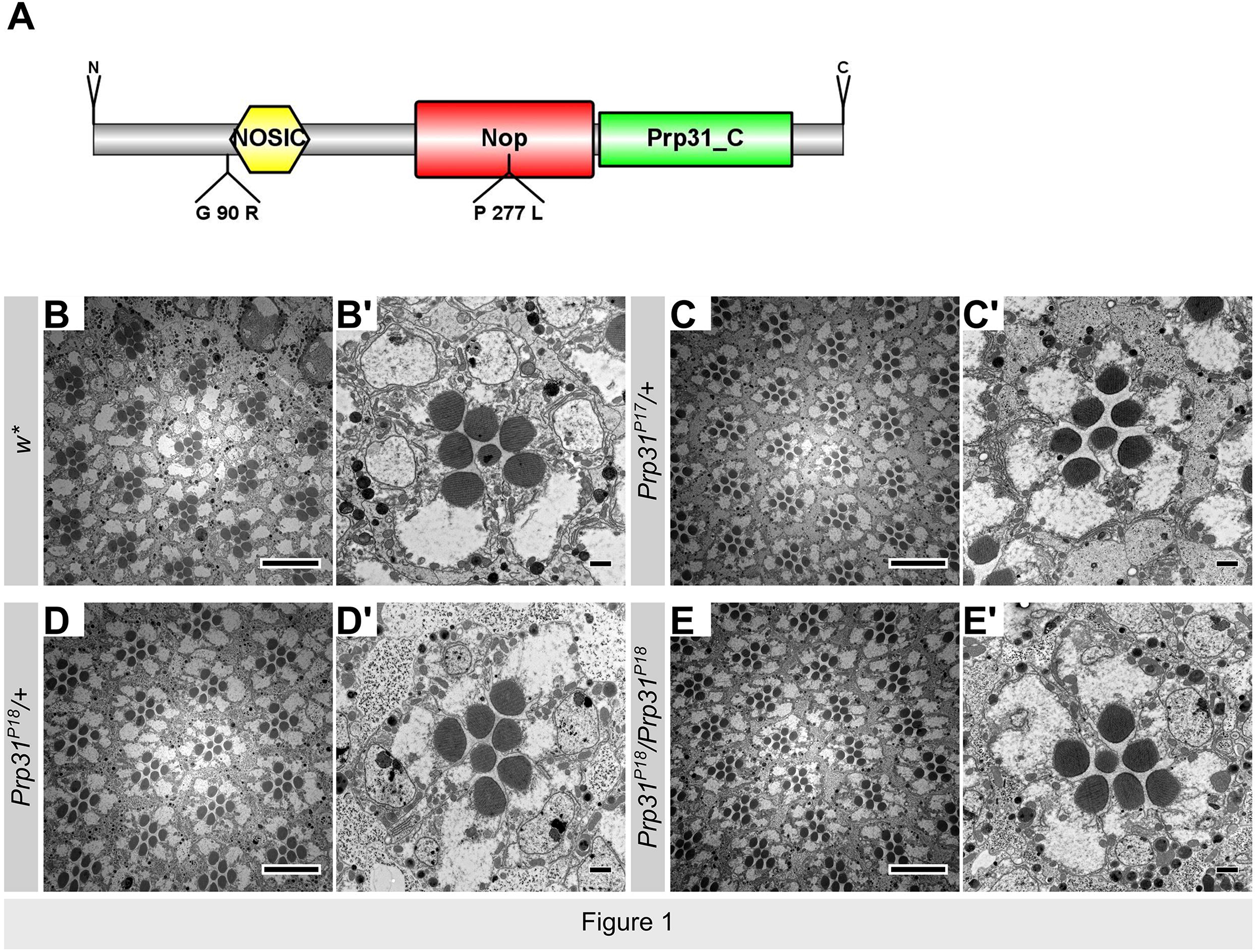
*Prp31* mutant flies have no gross morphological abnormalities at eclosion. (A) Schematic overview of the human Prpf31 protein. The figure is drawn to scale using IBS (Liu et al., 2015). Domains and TILLING mutations described here are indicated. (B-E’) are representative electron micrographs of 70nm sections of eyes of genetic control *w*\* (B, B’), *Prp31^P17^/*+ (C, C’), *Prp31^P18^/*+ (D, D’), and *Prp31^P18^/Prp31^P18^* (E, E’). Upon eclosion, flies were kept for two days under regular light conditions. B’, C’, D’ and E’ depict higher magnifications of one ommatidium of the respective overviews shown in B, C, D and E. Note that the number and stereotypic arrangement of photoreceptor cells within the mutant ommatidia are not affected. Scale bars represent 10 μm in B, C, D and E and 1 μm in B’, C’, D’ and E’.

Homo- and hemizygous *Prp3^P18^*, hemizygous *Prp3^P17^* flies as well as *Prp31^P17^*_/_*Prp31^P18^* transheterozygous flies kept under normal light/dark cycles have eyes of normal size. Histological sections revealed normal numbers of PRCs (distinguished by the number of rhabdomeres) per ommatidium and a normal stereotypic arrangement of PRCs (Fig. 1B-F). This indicates that the development of the retina was not affected by these mutations. However, PRCs of *Prp31^P17^*/+, *Prp31^P18^*/+ and *Prp31^P18^*/ *Prp31^P18^* flies showed severe signs of retinal degeneration when exposed to constant light for several days. The same phenotype was observed in PRCs of *Prp31^P17^* or *Prp31^P18^* hemizygous flies as well as in *Df(3L)Exel6262/*+ flies (Fig. 2B-C’ and data not shown). After seven days of light exposure, the majority of rhabdomeres of the six outer PRCs, R1-R6, were either completely gone or were strongly reduced in size, and many cell bodies exhibited the typical signs of degeneration, such as condensed chromatin and electron dense material with vesiculation and multivesicular bodies (Fig. 2B-C’; quantification in Fig. 2G). No significant differences were observed between the two alleles. R7 is preserved in most ommatidia. Strikingly, PRCs of transheterozygous *Prp31^P17^*/*Prp31^P18^* flies showed less degeneration in comparison to those of flies heterozygous for either of these alleles (Fig. 2E, E’). Many intact rhabdomeres were present, although some of them showed first signs of degeneration. Only few PRCs showed the dark staining typical for degenerating cells (quantification in Fig. 2F). In contrast to other fly models of retinal degeneration, for example *crb* (Pocha et al., 2011), *Prp31* mutant flies aged for 30 days under regular light/dark conditions did not show major signs of degeneration (data not shown). To summarise, the isolated mutant *Prp31* alleles reveal dominant, light-dependent retinal degeneration, similar as mutations in RP11 patients, and show intragenic complementation.

**Figure 2:**
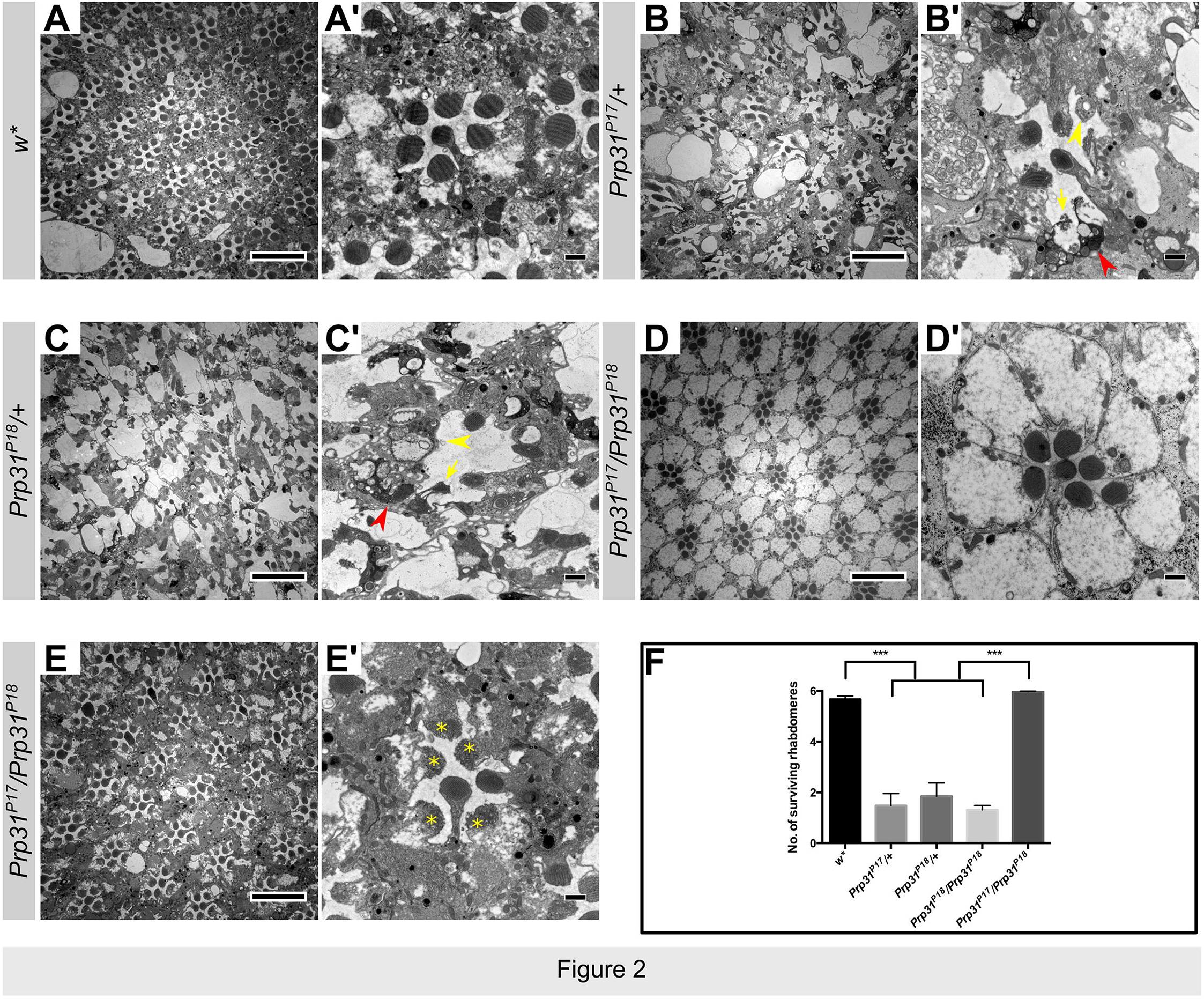
Flies heterozygous for *Prp31^P17^* or *Prp31^P18^* undergo light-dependent degeneration. (A-C’) are representative electron micrographs of 70nm sections of eyes of genetic control *w*\* (A, A’), *Prp31^P1^*^7^*/*+ (B, B’), *Prp31^P18^/+ (*C, C) exposed for 7 days to continuous, high intensity light. (D-E’) are representative electron micrographs *of Prp31^P17^/Prp31^P18^* eyes exposed to regular light conditions (12 hours light/12 hours dark; D, D’) and continuous high intensity light (E, E’). Note that degeneration under constant light exposure is more pronounced in *Prp31* heterozygotes (B, B’ and C, C’) than in transheterozyotes (E, E’). The most obvious features of degenerating PRCs are (i) absence (yellow arrowhead) or reduction (yellow arrow) of rhabdomeres of outer photoreceptor cells (R1-R6), (ii) accumulation of electron dense material with vacuolization (red arrowhead) as seen in B’ and C’. Transheterozygotes do not show any defect when kept under regular light conditions (D, D’), but exhibit loss of rhabdomeric integrity (yellow asterisks), which is considered as an early sign of degeneration, upon exposure to constant light (E, E’), Scale bars represent 10 μm in A, B, C, D and E and 1 μm in A’, B’, C’, D’ and E’. (F) Quantification of retinal degeneration. Bars represent mean ± s.e.m. for the number of surviving rhabdomeres of outer PRCs R1-R6 following high intensity, continuous light exposure. Significant differences are indicated by black horizontal bars. *** indicates significance at p<0.005 calculated using ANOVA and post-hoc Bonferroni correction.

### *Prp31^P18^* mutant flies exhibit increased oxidative stress signalling

Photoreceptors have an extraordinary oxygen consumption due to their high biosynthetic activity, which is required to continuously replenish the photosensitive apical membrane (Ng et al., 2015; Yu and Cringle, 2001). In addition, although PRCs are specialised for light reception to initiate phototransduction, light at the same time is a stress factor and induces increased production of reactive oxygen species (ROS) (German et al., 2015). Increased levels of cellular ROS, in turn, induce antioxidant responses, which include the expression of proteins against oxidative stress, e.g. superoxide dismutase (SOD) or glutathione Stransferase. Their activity can prevent the cell from the detrimental consequences of oxidative stress, such as increased lipid oxidation or damage of proteins and DNA (Tomanek, 2015). In photoreceptor cells, a failure of the antioxidant machinery to neutralise increased levels of ROS can lead to light-dependent retinal degeneration, for example in fly PRCs mutant for *crb* (Chartier et al., 2012).

This raised the question whether flies mutant for *Prp31* are subject to increased oxidative stress. To address this question, we analysed *Prp3^P18^* mutant flies that carried the *GstD-GFP* reporter transgene. This reporter expresses GFP under the control of upstream regulatory sequences of *glutathione S-transferase* (*gstD1*), one of the genes involved in detoxification, whose expression is activated by oxidative stress (Sykiotis and Bohmann, 2008). The expression of this reporter has been shown to correlate with ROS levels, as revealed by the ROS-sensitive dye Hydro-Cy3 in the midgut of adult flies stressed by feeding bacteria (Jones et al., 2013). GFP expression was determined by measuring fluorescence levels in lysates of flies kept for two days on standard food and of flies kept for three days on food containing 5% hydrogen peroxide (H_2_O_2_), an established oxidative stressor. Compared to control flies, *Prp31^P18^* heterozygous flies kept on normal food showed a 25% increase in *GstD-GFP* expression (Fig. 3A). This suggests that *Prp31^P18^* mutants are under increased levels of oxidative stress signalling already under standard/basal conditions. Upon exposure to 5% H_2_O_2_, lysates from control flies displayed an 18% increase in *GstD-GFP* expression, while lysates from *Prp31^P18^* mutant flies exhibited a 47% higher level of GFP compared to control flies kept on standard food (Fig. 3A). To corroborate these findings, GFP expression was examined in-situ by immunostaining of adult eye tissue. In control eyes, GstD-GFP expression was high in the pigment and cone cells. Interestingly, no GstD-GFP expression was detected in the photoreceptor cells themselves (Fig. 3B-B’). In eyes of *Prp31^P18^*/+ flies GstD-GFP levels were strongly increased in cone and pigment cells (Fig. 3C-C’). Taken together, these data show that *Prp31^P18^* mutant eyes exhibit increased oxidative stress signalling already under normal conditions, particularly in cone and pigment cells.

**Figure 3:**
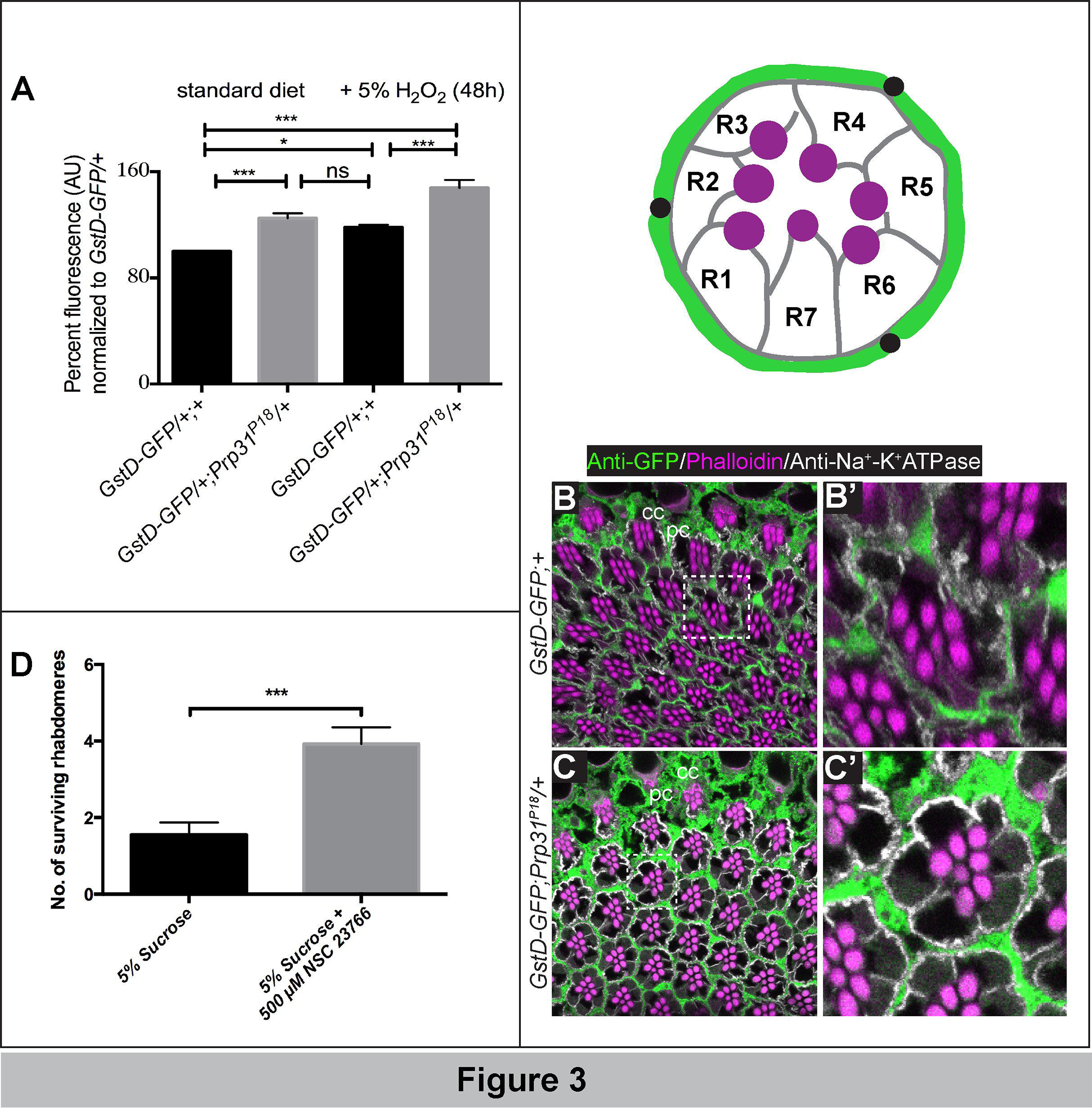
*Prp31^P18^* mutant flies exhibit increased oxidative stress signalling. A: Percent GFP fluorescence levels measured from lysates of adult flies of the indicated genotypes after normalization to fluorescence levels in control flies (*gstD-GFP/+)*, set at 100 (arbitrary units). Flies were fed on a standard yeast diet or on standard yeast food supplemented with 5% _H2O2._ The values were averaged from 5 biological replicates (standard food) and 2 biological replicates (5% H2O2). Comparisons are indicated by the black horizontal bars, * indicates significance at p<0.05, *** indicates significance at p<0.005 and ns indicates non-significance calculated using ANOVA followed by a post-hoc Bonferroni correction. Bars represent mean ± s.e.m. B-C’: Top is a cartoon of an ommatidium including the 7 photoreceptor cells (R1-R7) with their photosensitive organelle called the rhabdomere (magenta). Pigment cells (pc) lie adjacent to the photoreceptor cells. Bottom are representative 1μm confocal optical sections imaged from a 10μm section of an eye of GstD-GFP;+ (B, B’) or GstD-GFP; P*rp31^P18^* heterozygotes (C, C’). Sections have been labelled with anti-GFP (green; GstD activity), phalloidin (magenta; rhabdomeres) and anti-Na^+^-K^+^-ATPase (white; basal membranes of photoreceptor cells). Boxed outlines in B and C indicate the ommatidium shown at higher magnification in B’ and C’, respectively. In control eyes (B, B’), GFP is evident in cone cells (cc) and in the pigment cells (pc) surrounding each ommatidium. Gst-GFP expression is increased in *Prp31^P18^* mutants (C, C’). Note that in contrast to high GFP levels in support cells, no GFP was detected in photoreceptor cells. D: Quantification of photoreceptor degeneration of *Prp31^P18^* homozygous mutants fed with 5% sucrose or with 500 μM NSC 23766 in 5% sucrose. Graph represents mean ± s.e.m. of surviving rhabdomeres of outer PRCs R1-R6, measured after 7 days exposure to constant, high intensity light. Feeding with NSC 23766 significantly reduced degeneration. *** p<0.005 as determined by a two-tailed Mann-Whitney U test.

### Prevention of light-induced degeneration of *Prpf31* mutant photoreceptor cells

It is well documented that increased ROS levels in both vertebrate and invertebrate PRCs can lead to neuronal degeneration upon exposure to additional stress, such as light stress (Punzo et al., 2012). Therefore, we assumed that the observed increase in the antioxidant response in the retina of *Prp31* mutant flies is the result of increased ROS levels, and that these could be the cause for light-dependent retinal degeneration. To test this assumption, we experimentally blocked one of the major sources of cellular ROS production, mediated by the NOX (**N**ADPH **ox**idase) family (Bedard and Krause, 2007). One subunit of NOX is Rac1, a member of the small GTPase protein family, which is involved, besides cytoskeletal remodelling, in the generation of ROS (Hordijk, 2006). Previous reports suggested that light can stimulate Rac1 (Balasubramanian and Slepak, 2003; Belmonte et al., 2006), which in turn results in enhanced NADPH-oxidase activity and thus increased ROS levels. In mice, constitutively active Rac1 can promote photoreceptor degeneration (Song et al., 2016), while depletion of Rac1 in photoreceptor cells was able to protect the cells from photooxidative stress (Haruta et al., 2009; Song et al., 2016). Furthermore, inhibiting Rac1 activity in a fly model for RP12, caused by loss of *crb* function, reduced ROS production and prevented light-dependent retinal degeneration (Chartier et al., 2012). To test whether NOX is involved in light-dependent retinal degeneration in *Prp31^P18^* homozygous mutants, we blocked Rac1 activation by feeding flies for two days before and during exposure to light with NSC23766, a selective inhibitor of the Rac1-GEF interaction and hence of Rac1 activation (Nassar et al., 2006). After light-exposure, treated animals exhibited a strongly reduced number of apoptotic PRCs (Fig. 3D). From these results we conclude, that increased NADPH-oxidase activity, which is likely followed by increased accumulation of ROS, is a major trigger of light-dependent PRC degeneration in *Prp31^P18^* homozygous mutant flies.

It has been suggested that retinal degeneration in *Prpf31* mutant mice is caused by accumulation of truncated, and hence toxic rhodopsin proteins, which is generated as a result of improper splicing of the rhodopsin pre-mRNA (Yuan et al., 2005). To find out whether rhodopsin accumulation may contribute to light-dependent degeneration in *Prp31* mutant flies, we raised larvae in a food lacking vitamin A, the precursor for retinal, the cofactor covalently bound to opsin. This treatment was shown to prevent light-dependent retinal degeneration in *crb* mutant PRCs (Johnson et al., 2002). As shown in Fig. 4, lack of dietary vitamin A strongly suppressed light-dependent degeneration of PRCs in *Prp31* mutant flies.

**Figure 4:**
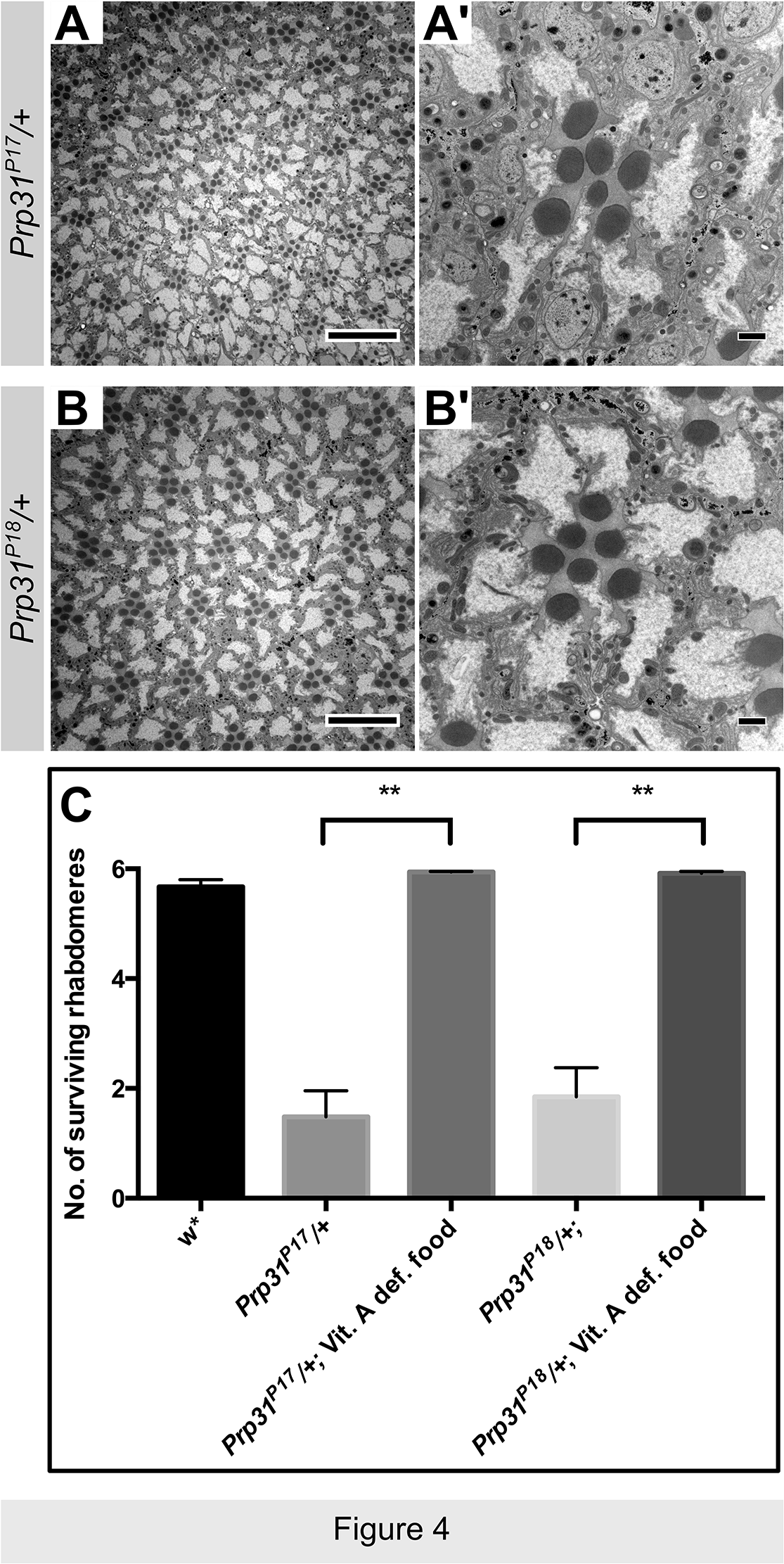
Raising *Prp31* mutant animals in vitamin A-depleted food rescues light-dependent degeneration. (A-B’) are representative electron micrographs of 70nm sections of eyes of *Prp31^P1 7^/*+ (A, A’) and *Prp31^P18^/*+ (B, B’) raised on vitamin A deficient food and placed in constant, high intensity light for seven days. Rhabdomeres are smaller due to vitamin A deficiency, but overall there are no obvious signs of retinal degeneration (compare to Fig. 2C, C’). Scale bars represent 10 μm in A and B and 1 μm in A’ and B’ (C) Quantification of photoreceptor degeneration in *Prp31* heterozygous mutants raised with normal and with vitamin A-depleted food. Bars represent mean ± s.e.m. of surviving rhabdomeres of outer PRCs R1-R6 in *Prp31^P17^/*+ and *Prp31^P18^/*+ flies raised and kept on vitamin A deficient food after seven days of exposure to constant light. ** indicates significance at p<0.01 calculated using a two-tailed Mann-Whitney U test.

### *Prp31* mutants reveal mis-regulation of genes involved in oxidation-reduction

In mouse and zebrafish, PRC degeneration due to loss of *PRPF31* has been associated with a general reduction in biosynthetic activity as well as with impaired splicing of PRC-specific mRNAs, such as *rhodopsin*, RDS (Peripherin) or Fascin (FSCN2) mRNA (Linder et al., 2011; Mordes et al., 2006; Tanackovic et al., 2011; Yin et al., 2011; Yuan et al., 2005). To get a deeper insight into the mechanisms by which *Drosophila Prp31* prevents retinal degeneration, we performed whole transcriptome analysis from RNA isolated from heads of *Prp31^P18^*/*Prp31^P18^* mutant and control (*w*) flies (2 days old, kept under normal light/dark conditions). For each genotype three biological replicates were analysed. Overall, 115 genes were significantly (q-value cut-off 0.01) and differentially expressed (2-fold change) in *Prp31^P18^* homozygous fly heads compared to *w* control flies. Of these, 53 were up-regulated and 62 were down-regulated (Suppl. Table S1, S2). Differentially expressed genes were categorized using the Gene Ontology (GO)/PANTHER classification system (Mi et al., 2013; Thomas et al., 2003), based on the predicted protein class of their gene products. Within the groups of up- and down-regulated genes, the three most prominent categories comprised genes with hydrolase, oxidoreductase, and transporter activities (Fig. 5A, B).

**Figure 5:**
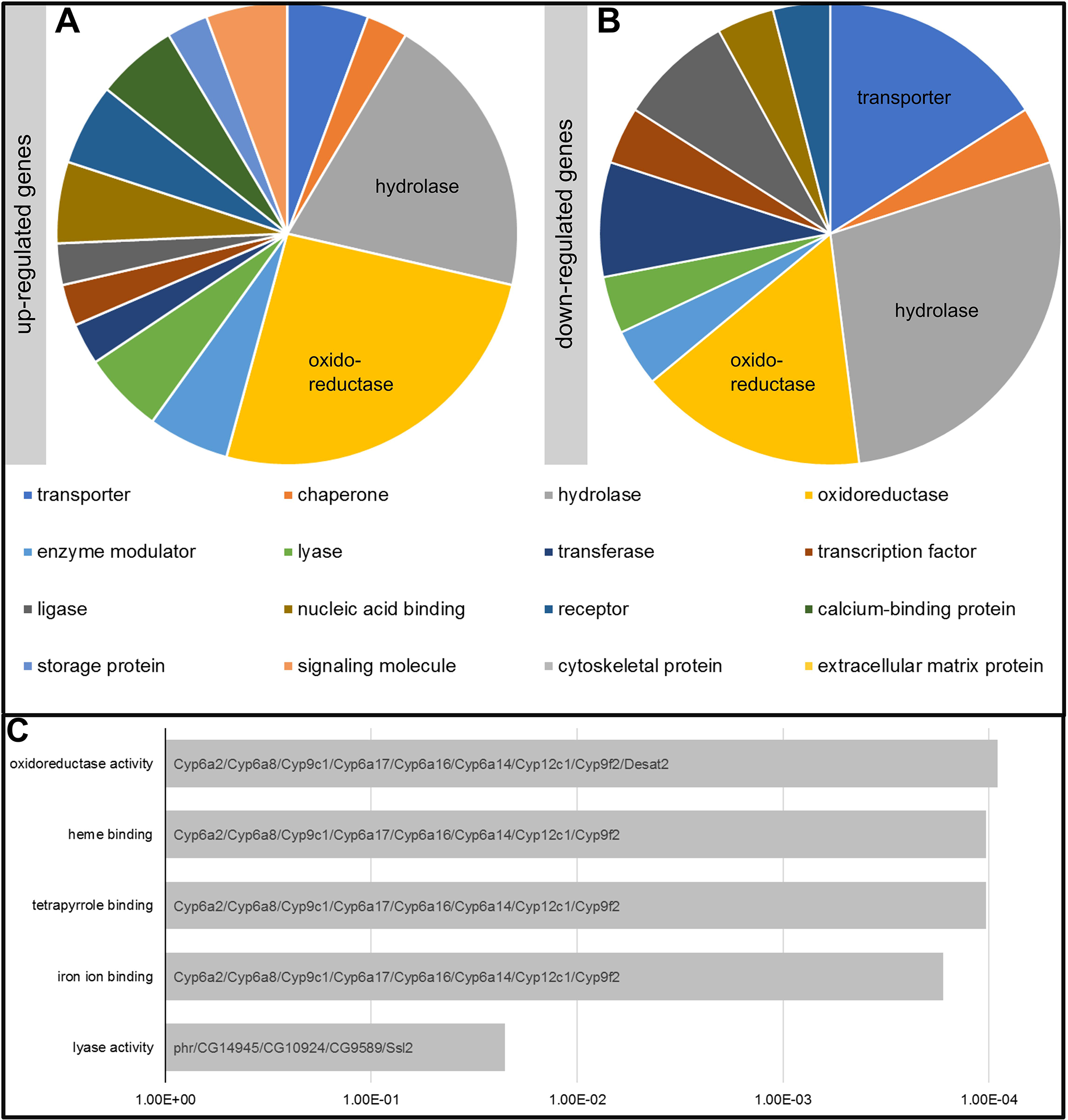
Classification of Up- and Down-regulated genes in *Prp31^P18^* fly heads. (A-B) Pie charts showing the classification of the 52 genes with significantly increased (A) and the 63 genes with significantly decreased (B) expression in mutant *Prp31^P18^/Prp31^P18^* heads, compared to control heads using the Gene Ontology/PANTHER classification system, based on the predicted protein classes for their gene products. Colors represent the different categories in the pie charts. (C) Bars show the significantly enriched GO terms for all genes that are differentially expressed in the mutant, compared to the genetic control, classified according to the GO category “molecular function”. The Y-axis indicates the sub-categories, the X-axis indicates the significance of the enrichment in GO terms compared to genes in total. Gene symbols of the genes falling into the category are stated in the bars.

We then focused on the 53 genes with increased expression in the mutants (Table S1). The largest group of those that match a GO term (n=9) fall into the GO term: oxidoreductase (Fig. 5A). This group includes genes of the Cytochrome-P450 family as well as the gene *cinnabar* (*cn*), which encodes a monoxygenase with a predicted NAD(P)H oxidase activity, and plays a role in the pigment biosynthetic pathway in the eye. We also observed an up-regulation of *w*, encoding a transporter, and *Rh6*, which encodes an opsin expressed in a subset of photoreceptors. Amongst the 62 genes with decreased expression in the mutant (Table S2) the three most abundant classes matching a GO term comprised genes encoding hydrolases, oxidoreductases, and transporters (Fig. 5B). The group of hydrolases contains a chitinase (*Cht3*). Relevant for eye function are genes encoding transporters, including the gene *scarlet* (*st*), which is known to be involved in pigment formation in the retina.

Since we observed overlapping categories in genes with increased and decreased expression in the mutants, we evaluated all 115 differentially expressed genes by applying statistical enrichment tests. Through this we identified enrichment for oxidoreductase activity, heme binding, tetrapyrrole binding, and lyase activity (Fig. 5C), corroborating our qualitative results. It should be noted that with the exception of *Desat2* in the first group (oxido-reductase activity) the first four classes comprise the same 8 members of the Cytochrome-P450 (*Cyp*) gene family.

### *Prp31* mutants exhibit increased intron retention

Since *Prp31* encodes a splicing factor, we asked whether a mutation in this gene impairs splicing on a genome-wide level, as has been described in zebrafish eyes with reduced *Prpf31* activity (Linder et al., 2011), in *Drosophila* embryos mutant for *Prp19* (Sauerwald et al., 2017), and in lymphoblasts derived from patients mutant for *PRPF3*, *PRPF8* or *PRPF31*(Tanackovic et al., 2011). Our data reveal that from a total of 49345 introns encoded in the genome, present in 13917 genes, 131 were significantly retained (FDR<0.1) in the mutant transcriptome, representing 127 genes (Table S3). To further evaluate the genes showing intron retention, we applied statistical enrichment tests. Strikingly, introns derived from genes falling into the categories of synaptic vesicle cycle/localization/exocytosis/transmission were particularly retained in the mutant (Fig. 6).

**Figure 6:**
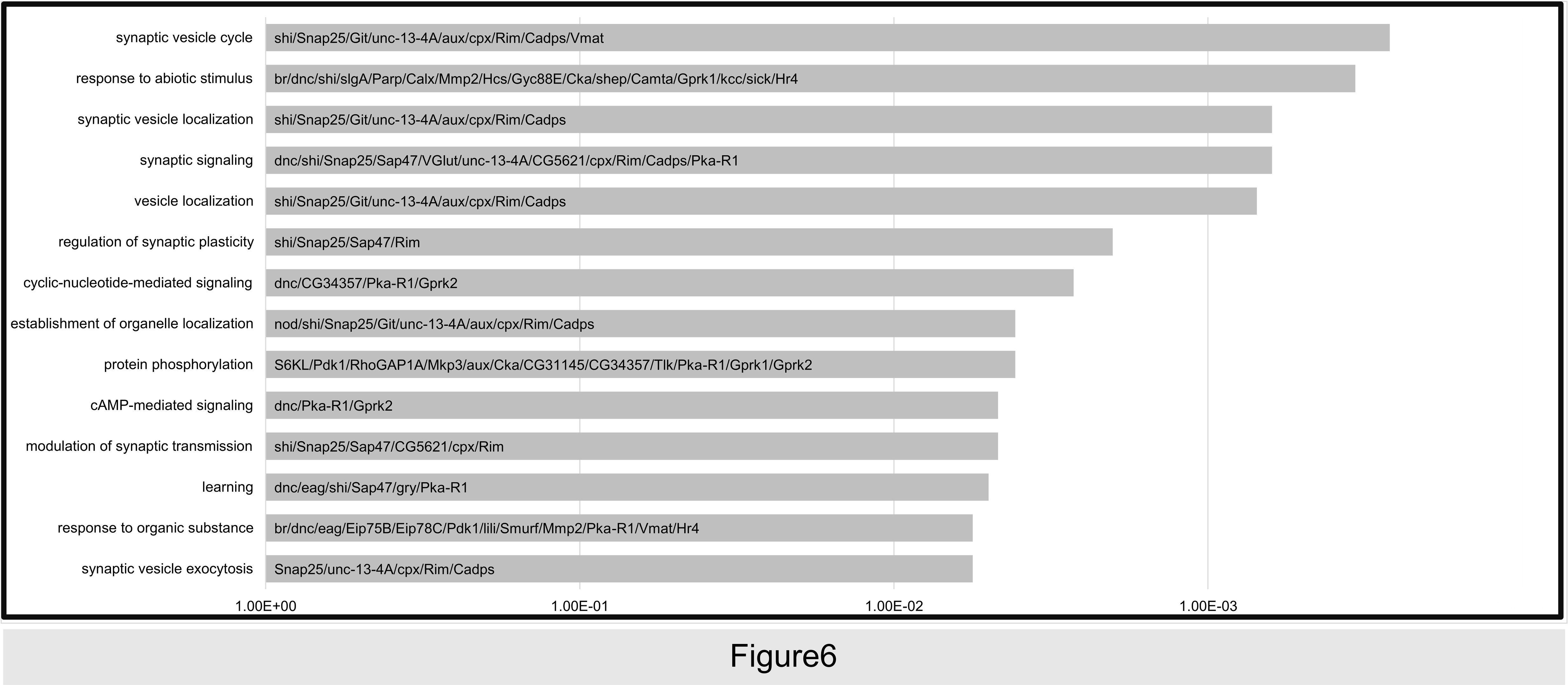
Classification of retained introns in *Prp31^P18^* mutants. Bars show the significantly enriched GO terms for genes that display intron retention in the mutant, compared to the genetic control, classified according to the GO category “biological process” (p-value < 1.12E-04). The Y-axis indicates the sub-categories, the X-axis indicates the significance of the enrichment in GO terms compared to genes in total. Gene symbols of the genes falling into the individual categories are listed.

Taken together, we established a fly model for RP11, a retinal disease caused by mutations in the highly conserved splicing factor PRPF31. We show that mutations in *Drosophila Prp31* induce light-dependent retinal degeneration, thus mimicking the symptoms of the human disease. We further uncovered major dysregulation of the transcriptome of mutant fly heads. The nature of the mis-regulated genes as well as the result of rescuing the mutant phenotype let us to conclude that light-dependent PRC degeneration of *Prp31* mutant eyes is of multifactorial origin, including increased oxidative stress, defects in rhodopsin metabolism, and splicing defects in genes involved synaptic transmission.

## Discussion

Here we present a fly model for RP11, an autosomal-dominant human disease leading to blindness, which is caused by mutations in the splicing regulator PRPF31. Our results reveal that, similar as in humans, mutations in the *Drosophila* orthologue *Prp31* lead to PRC degeneration under light stress, thus mimicking major features of RP11-associated symptoms. Similar as in human, mutations in *Drosophila Prp31* lead to retinal degeneration when heterozygous. This is in stark contrast to a mouse heterozygous for *Prpf31*, which did not show any signs of retinal degeneration (Bujakowska et al., 2009). Different studies showed late-onset defects in the retinal pigment epithelium of *Prpf31* mutant mice (Farkas et al., 2014; Graziotto et al., 2011). Most of the mutations in human *PRPF31* linked with RP11 are associated with reduced *PRPF31* mRNA, suggesting that these are loss-of-function alleles (Rio Frio et al., 2008; Ruzickova and Stanek, 2016).

Data presented here let us to conclude that the two missense mutations mapped in *Prp31^P17^* and *Prp31^P18^* represent hypomorphic conditions, which reduce, but do not abolish the function of the protein. First, unlike in humans, the two *Drosophila* alleles characterized here are hemizygous and homozygous (in the case of *Prp31^P18^*) viable and fertile. Second, *Prp31^P17^*/*Prp31^P18^* flies showed intragenic complementation and exhibited only a mild degenerative phenotype. Intragenic (interallelic) complementation is a rare event, and is often explained by the fact that two or more defective proteins can form functional multimers, if their mutations reside in different domains. This has been shown, for example, for mutations affecting *Drosophila* Dynein (Gepner et al., 1996) or Posterior sex combs (Psc), a member of the homeotic Polycomb Group (PcG) proteins involved in epigenetic silencing (Wu and Howe, 1995). In fact, the mutations in the two established *Prp31* fly lines reside in different parts of the protein, namely N-terminal to the NOSIC domain in *Prp31^P17^* (G90R) and in the Nop domain in *Prp31^P18^* (P277L) (see Fig. 1A). However, it still remains to be analysed whether Prp31 proteins dimerize. As shown in yeast, Prp31 is a component of the spliceosomal U4/U6 di-SNP, which contains, beside the base-paired U4 and U6 snRNAs, more than 10 other proteins, including Prp3 and Prp4. In this complex, Prp31 is required to stabilize a U4/U6 snRNA junction, which in turn is required for binding of Prp3/4 (Hardin et al., 2015). In human PRPF31, the Nop domain is involved in an essential step in the formation of the U4/U6-U5 tri-snRNP by building a complex of the U4 snRNA and a 15.5K protein. Consistent with this, many mutations in human PRPF31, which are linked to RP11, have been mapped to the Nop domain. Mutations in amino acid H270 in the Nop domain of human PRPF31 results in its reduced affinity to a complex formed by a stem-loop structure of the U4 snRNA and the 15.5K protein (Liu et al., 2007; Schultz et al., 2006). Interestingly, the mutated amino acid residue in *Drosophila Prp31^P18^* (P277L) lies next to H278, which corresponds to amino acid H270 in the human protein. Therefore, it is tempting to speculate that the *Drosophila* P277L mutation could similarly weaken, but not abolish the corresponding interaction of the mutant Prp31 protein in flies, thus explaining the hypomorphic nature of this allele. Finally, PRC-specific RNAi-mediated knock-down of *Prp31* induced a similar phenotype as the one observed in *Prp31* heterozygous animals [data not shown and (Ray et al., 2010)]. The defects on eye morphogenesis detected upon more widespread knock-down of *Prp31* (Ray et al., 2010) indicates that *Prp31* may be important in other tissues as well, supporting our assumption that *Prp31^P17^* and *Prp31^P18^* are hypomorphic alleles. Further experiments are required to determine the functional consequence of the molecular lesions. Preliminary results make it unlikely that impaired nuclear localisation of the mutant proteins is responsible for the reduced function: the mutant proteins localise to nuclear speckles when expressed in HeLa cells (data not shown), and hence behave similar as the wild-type protein (Makarova et al., 2002).

Our results show that *Prp31* heterozygous flies undergo retinal degeneration, a phenotype with striking similarity to human RP11 patients. This now allows to further unveil the cause of the aetiology of the disease in a model organism, which is easily accessible to genetic manipulations and molecular studies (Ugur et al., 2016). Our results propose a multifactorial genesis of *Prp31*-linked retinal degeneration in flies. i) We suggest that increased accumulation of intracellular rhodopsin may contribute to the degeneration in *Prp31* mutant retinas. Immunostainings of retinae of hetero-, as well as homozygous *Prp31^P18^* flies showed an accumulation of Rh1 in the photoreceptor cell body in comparison to *w^1118^* control flies (data not shown). Accumulation of misfolded rhodopsin in the ER due to dominant mutations in the gene which impair the maturation of the protein, has been described to cause an overproduction of ER cisternae and eventually leads to degeneration (Colley et al., 1995). Interestingly, mis-localisation of rhodopsin in human PRCs to sites other than the outer segment is a common characteristic of various forms of RP and is considered to contribute to the pathological severity (Hollingsworth and Gross, 2012). Light-dependent PRC degeneration in *Prp31*-mutant flies could be prevented by raising mutant animals with food that lacks vitamin A, the precursor for retinal. This treatment reduces the amount of rhodopsin to about 3% of its normal content (Nichols and Pak, 1985), and thus strongly reduces rhodopsin accumulation in the cell body.

ii) Our data further suggest that besides accumulation of intracellular rhodopsin increased oxidative stress contributes to light-dependent PRC degeneration in *Prp31* mutant flies, since the degree of degeneration could be reduced by feeding flies with NSC23766, a known inhibitor of NADPH oxidase (NOX) activity. NADPH oxidases are multi-subunit enzyme complexes, comprising two membrane-bound and three cytoplasmic components, and can be found in the plasma-membrane of many cell types. One of the cytoplasmic components is the small GTPase Rac1 (Hordijk, 2006). Recruitment of the cytoplasmic components to the membrane activates the complex, resulting in transfer of electrons from NADPH to molecular oxygen, thus producing superoxide. NSC23766 selectively inhibits the interaction between Rac1 and Rac1-specific guanine nucleotide exchange factors (GEFs), thereby preventing the activation of Rac1 and hence NOX activity (Katsuyama, 2010; Rastogi et al., 2016). The suppression of retinal degeneration in *Prp31* mutant flies by NSC23766 suggests that increased ROS production is causally related to degeneration. This assumption is supported by the observation that mutant flies show enhanced expression of GstD1-GFP, a reporter that serves as a proxy of ROS levels (Sykiotis and Bohmann, 2008).

Data obtained from transcriptome analysis of *Prp31* mutant heads strongly support the conclusion that oxidative stress contributes to the mutant phenotype. The genes most highly upregulated in heads of *Prp31* mutant flies match GO terms that are known to be upregulated upon oxidative stress (Girardot et al., 2006; Landis et al., 2004). These are, for example genes in involved in oxidation/reduction processes, e.g. *desat2*, *Cytochrome P450-6a17* (*Cyp6a17*) and *Cyp9c1*. Some of the genes, which are downregulated in *Prp31* mutant fly heads are those involved in polysaccharide/chitin metabolisms, e. g. *Cht3*, encoding a chitinase, and *Tweedle E* (*TwdlE*), a cuticular protein (Cornman, 2009; Guan et al., 2006; Karouzou et al., 2007). Genes from this category were shown to be highly up-regulated in flies exposed to mild stress by increased atmospheric pressure (hyperbaric normoxia) (Yu et al., 2016). This stress has been suggested to induce cytoprotective responses, aimed to protect cells or organisms against the detrimental effects of stressors, such as senescence (Oh et al., 2008). Chitin oligosaccharides comprise a major constituent of the insect cuticle and have been suggested to function as antioxidants by scavenging ROS (Ngo and Kim, 2014) and to induce the immune response (Li et al., 2013). The downregulation of these genes in *Prp31* mutant heads points to a reduced cytoprotective response and hence may contribute to the detrimental effects of light stress.

Taken together, the retina of *Prp31* mutant flies exhibits increased oxidative stress response, suggesting that upon additional stress, provided by constant light exposure, the antioxidant response machinery is no longer able to prevent the damage induced by high levels of ROS. This may lead, among others, to oxidation of lipids, which are major constituents of the photosensitive organelle, the outer segments in vertebrates and the rhabdomeres in flies. Oxidised lipids may contribute to PRC degeneration, as shown for Age-related Macular Degeneration (AMD) (Handa et al., 2017).

iii) Data from the transcriptome analysis further reveal that impaired *Prp31* function results in increased intron retention, suggesting defects in splicing. It is worth mentioning that with the exception of *Adh*, impaired splicing did not overlap with the down-regulated genes. From this we conclude that transcript down-regulation largely does not reflect nonsense-mediated mRNA decay, which often occurs in order to prevent translation of intron-containing mRNAs into non-functional and potentially detrimental proteins (Yap and Makeyev, 2013). Interestingly, genes with the most significant intron retention are those involved in regulating the function, localisation and maturation of synaptic vesicles. Whether this results in defective synaptic transmission, remains to be elucidated. Previous work has shown that mutations in genes required for proper synapse organization in PRCs may eventually result in retinal degeneration, e. g mutations in *Drosophila Lin-7/veli* (Soukup et al., 2013). Similarly, mislocalization of pre- and postsynaptic proteins, as observed in *rd1* and *rd10* mutant mice [reviewed in (Soto and Kerschensteiner, 2015)], or structural abnormalities in PRC synapses as observed in *tulp* mutant mice (Grossman et al., 2009) precede PRC degeneration.

Taken together, using the fly as a genetic model revealed a multifactorial genesis of light-dependent retinal degeneration of *Prp31* mutant flies. Impaired *Prp31* function impacts on various cellular processes and results in increased oxidative stress, defective rhodopsin transport/maturation and intron retention, predominantly in transcripts of genes involved in synapse formation and function. To what extend these dysfunctions influence each other has to be elucidated.

## Materials and Methods

### Fly maintenance and genetics

Flies and crosses were maintained at 25°C, on standard yeast-cornmeal-agar food, under 12 hours of light/12 hours of darkness, unless specified. Flies were placed for a total of 7 days under these conditions. Genetic control for all experiments was *white* (*w*\*). Deficiency lines used here were obtained from the Bloomington Stock Centre and included *Df(3L)Exel6262* (Parks et al., 2004)*, Df(3L)ED217and Df(3L)ED218* (Ryder et al., 2007). *GstD-GFP* flies (Sykiotis and Bohmann, 2008) (gift from D. Bohman) were used as an indicator of oxidative stress signalling by combining into the *Prp31^18^* genetic background or into the genetic control (*w*\*). For the light-stress experimental paradigm, flies were kept at 25°C for 7 days in a special incubator designed to have high intensity (1200-1300 lux), continuous light exposure (Johnson et al., 2002)

### Anti-oxidant feeding

Female flies were collected immediately upon eclosion and divided into groups maintained on regular food, supplemented with a Whatman^®^ filter paper (GE Healthcare) soaked in 5% sucrose (control) or with 500 μM NSC 23766 (Cayman Chemical). Food and paper were changed every second day.

### Vitamin A depletion

For vitamin A depletion experiments, flies were raised from embryonic stages until adulthood and subsequently maintained on carotenoid free food (10% dry yeast, 10% sucrose, 0.02% cholesterol, and 2% agar) as described (Pocha et al., 2011).

### Hydrogen Peroxide exposure

Flies were raised on standard yeast-food and upon eclosion, were transferred in groups of 10 onto standard food or food supplemented with 5% H2O2 (Sigma-Aldrich, Germany). After two days under 12h light and 12h dark conditions, 4-6 flies/genotype were used for further analyses

### Quantification of *GstD-GFP* following H_2_O_2_ feeding

Flies were lysed in 200μl of phosphate buffer (pH 7.4-7.6) with 0.1% Tween-20 on ice. Lysates were centrifuged at 15.000 rpm for 10 minutes. Of this, 25μl was used to estimate protein content by BCA assay and 150μl was used for fluorescence measurements using a Plate reader (Perkin Elmer Envision) and 485nm excitation & 590-10nm Emission filters. To calculate percent change, fluorescence values were normalized after arbitrarily setting values of control flies (*gstDGFP/+)* at 100. Data from 5 biological replicates (standard food) and 2 biological replicates (5% H_2_O_2_) were used for the analyses. ANOVA and post-hoc Bonferroni test was used to compare different samples.

### Isolation of *Prp31* alleles by TILLING

To isolate point mutations in the *Prp31* locus (FlyBase ID: FBgn0036487) a library, of 2.400 fly lines with isogenized third chromosomes which potentially carry point mutations caused by EMS treatment, was screened. Our approach targeted exon 1-3 of the Prp31 locus containing two thirds (67%) of the coding sequence and including several predicted functional domains (the NOSIC (IPRO012976), the Nop (IPRO002687) and parts of the Prp31_C terminal (IPRO019175) domain), making use of two different PCR amplicons. A nested PCR approach was used, where the inner primers contain universal M13 tails that serve as primer binding sites of the Sanger sequencing reaction:

- amplicon1 (covers exon 1 and 2), outer primer, forward: TTCAATGAACCGCATGG, reverse: GTCGATCTTTGCCTTCTCC, inner / nested primer, forward: TGTAAAACGA CGGCCAGT-AGCAACGGTCACTTCAATTC, reverse: AGGAAACAGCTATGACCAT-GAAAGGGAATGGGATTCAG);
- amplicon 2 (covers exon 3), outer primer, forward: ATCGTGGGTGAAATCGAG, reverse: TGGTCTTCTCATCCACCTG, inner / nested primer, forward: TGTAAAACGA CGGCCAGT-AAGCTGCAGGCTATTCTCAC, reverse: AGGAAACAGCTATGACCAT-TAGGCATCCTCTTCGATCTG.

PCR-reactions were performed in 10 μl volume and with an annealing temperature of 57 °C, in 384 well format, making use of automated liquid handling tools. PCR fragments were sequenced by Sanger sequencing optimized for amplicon re-sequencing in a large-scale format (Winkler et al., 2011; Winkler et al., 2005). Primary hits, resembling sequence variants, which upon translation result in potential nonsense and missense mutations or affect a predicted splice site, were verified in an independent PCR amplification and Sanger sequencing reaction.

### Transmission electron microscopy

Fixation of adult eyes, semi-thin sections and ultra-thin sections for transmission electron microscopy was performed as described (Mishra and Knust, 2013). 2 μm semi-thin sections were stained with a toluidine blue 1% / sodium tetraborate dehydrate 0.5% solution and imaged with AxioImager.Z1 (Zeiss, Germany) with an AxioCamMRm and the AxioVision software (Release 4.7). 70nm ultrathin sections were imaged using a Morgagni 268 TEM (100kV) electron microscope (FEI Company), and images were taken using a Side-entry Morada CCD Camera (11 Megapixels, Olympus).

### Quantification of Degeneration

Quantification of degeneration was performed as described in (Bulgakova et al., 2010). Briefly, from the semi-thin sections, the number of outer photoreceptor cells (R1-R6) with clearly detectable rhabdomeres in each ommatidium were recorded as surviving rhabdomeres. In each section, 50-60 ommatidia were counted and for each genotype, 6 eyes from different individuals were analysed, unless indicated. ANOVA and post-hoc Bonferroni correction and two-tailed Mann-Whitney U test, respectively, were used to compare different distributions of photoreceptor cell survival.

### Cryosections of *Drosophila* eyes

Adult eyes were dissected and fixed in 4% formaldehyde. Following sucrose treatment and embedding of the tissues in Richard-Allan Scientific NEG50^TM^ (Thermo Fisher Scientific, UK) tissue embedding medium, tissues were cryosectioned at 10μm thickness at −21°C. Sections were air-dried and then subjected to immunostaining as described previously (Spannl et al., 2017). Antibodies used were rabbit anti-GFP (1:500; A11122; Thermo Fisher Scientific, UK) and mouse anti-Na^+^-K^+^-ATPase (1:100; a5; Developmental Studies Hybridoma Bank, University of Iowa, USA). Alexa-Flour conjugated secondary antibodies (Thermo Fisher Scientific, UK) were used. F-actin was visualised with Alexa-Fluor-555–phalloidin (Thermo Fisher Scientific, UK). Images were taken on Olympus FV100 and processed using ImageJ/Fiji, Adobe Photoshop CS5.1 and Adobe Illustrator CS3 for image assembly.

### RNA extraction

Whole RNA was extracted from fly heads (2 days old) with the RNeasy Mini Kit (Qiagen) according to the manufacturer’s instructions. Each biological replicate consisted of 50 heads. A total of 3 biological replicates were used for further analyses.

### RNA-seq and analyses

RNA-Seq was carried out in triplicate by the Deep Sequencing Group SFB 655 at Biotechnology Center, TU Dresden, with 76-bp single read sequencing on the Illumina HiSeq 2500. An average of 28.85 million reads per sample was obtained. Data analysis was carried out by the Scientific Computing Facility (MPI-CBG) by mapping to the *Drosophila* genome (BDGP6, Ensembl v81) using STAR (v2.5.1b). The differential expression analysis was performed using DESeq2 using a lfcThreshold parameter of 1.0 and Independent Hypothesis Weighting (Love et al., 2014). 115 differentially expressed genes were obtained with a q-value cutoff of 0.01. Detection of differentially spliced genes was carried out using the DEXSeq Bioconductor package (Anders et al., 2012) with a discovery rate (FDR) threshold of 0.1. To test for intron retention as well as for differential exon usage, the DEXSeq model used to estimate exon abundance was complemented with intron regions (similar to the method presented by (Sauerwald et al., 2017)). clusterProfiler (Yu et al., 2012) was used to identify enrichment on a 0.05 q-value level for biological processes and pathways associated with the up- and down-regulated genes and for genes whose introns are differentially retained. The transcriptome data have been submitted to GEO. The accession number is GSE99665.

### Figure panel preparation

All figure panels were assembled using Adobe Illustrator CS3 and Inkscape. Statistical analyses and graphs were generated using GraphPad Prism (GraphPad Software, Inc, USA) and Microsoft Excel. For protein sequence visualization, Illustrator of Biological Sequences (IBS; (Liu et al., 2015)) software package was used.

---

**Supplementary Figure 1**: Amino acid sequence comparison of *Drosophila* Prp31 (upper) and human PRPF31 protein (lower). The NOSIC (yellow), Nop (red) and Prp31_C specific (green) domains are indicated as described on UniProt and Pfam websites. TILLING mutations are indicated in magenta. Asterisks (*) indicate fully conserved amino acids, colons (:) indicate groups of amino acids of strongly similar properties and periods (.) indicate amino acids with weakly similar properties. Alignment was made using ClustalO 1.2.3.

